# Benfotiamine reduces pathology and improves muscle function in *mdx* mice

**DOI:** 10.1101/288621

**Authors:** Keryn G. Woodman, Chantal A. Coles, Su L Toulson, Elizabeth M. Gibbs, Matthew Knight, Matthew McDonagh, Rachelle H. Crosbie-Watson, Shireen R. Lamandé, Jason D. White

## Abstract

Duchenne Muscular Dystrophy (DMD) is a progressive and fatal neuromuscular disease which arises from mutations in the dystrophin gene (*DMD*) that result in the absence or severe reduction of the cytoskeletal protein dystrophin. In addition to the primary dystrophin defect, secondary processes such as inflammation, calcium influx, dysregulated autophagy and fibrosis exacerbate dystrophic pathology and thus increase disease progression. While therapies to restore dystrophin deficiency are being developed, strategies which target these secondary processes could be of benefit to patients. Benfotiamine is a lipid soluble precursor to thiamine that can reduce secondary processes such as inflammation and oxidative stress in diabetic patients. As such we tested it in the *mdx* mouse model of DMD and found that benfotiamine reduced multiple markers of dystrophic pathology and improved grip strength. In addition, members of the utrophin and dystrophin glycoprotein complexes were significantly increased at the sarcolemma which could improve cell adhesion. We also demonstrated that benfotiamine treatment lowered the expression of macrophage markers and pro-inflammatory cytokines suggesting that benfotiamine is reducing dystrophic pathology by acting on inflammatory processes.

## Introduction

The muscular dystrophies are a group of genetic neuromuscular disorders that result in the progressive deterioration of skeletal muscle. Of these muscular dystrophies, Duchenne muscular dystrophy (DMD) is the most common affecting 1 out of every 5000 male births (1). This devastating condition results from mutations in the dystrophin gene leading to absence or severe reduction in dystrophin at the muscle plasma membrane (2). Dystrophin is a critical component of a large complex known as the dystrophin associated glycoprotein complex (DGC), present on the plasma membrane of the myofibre (3, 4). Dystrophin stabilises cells by linking actin filaments, intermediate filaments and microtubules to transmembrane complexes (5). Loss of dystrophin, and in most cases loss or reduction of the DGC, leads to membrane instability, increased susceptibility to mechanical stress and finally, degeneration of myofibres (6). Skeletal muscle possesses an innate ability to regenerate in response to injury but in DMD this is compromised over time due to the chronic nature of the damage and persistence of inflammatory cells (7). While the absence of dystrophin is the primary defect in DMD, it is becoming increasingly apparent that several secondary processes contribute to disease progression and associated muscle wasting in DMD. These include the activation of calcium influx, secretion of pro-inflammatory and pro-fibrotic mediators and defects in the clearance of damaged organelles through autophagy (8).

Normal muscle repair is a complex and highly regulated biological process. Following acute muscle injury resident macrophages and T lymphocytes are activated and other immune cells, including neutrophils, migrate to the injured tissue. The early immune response is driven by interferon and TNFα stimulation of M1 (classically activated) macrophages, which produce pro-inflammatory cytokines such as TNFα and IL-1β and remove the damaged tissue. After 1-3 days, the dominant macrophages become M2c which release anti-inflammatory cytokines, IL-10 and TGFβ, to deactivate M1 macrophages and promote repair (9-11). M2a macrophages (alternatively activated) are abundant later in the repair process and produce pro-fibrotic molecules such TGFβ. Macrophage-produced pro-inflammatory cytokines activate satellite cells and promote myoblast proliferation, while anti-inflammatory cytokines stimulate myoblast differentiation and fusion. Fibroblasts migrate into damaged muscle and are stimulated by TGFβ and pro-inflammatory cytokines to produce and remodel the extracellular matrix (ECM). While ECM deposition is needed for efficient repair, precise regulation is crucial and dysregulation leads to fibrosis (12).

Muscle repair thus involves the co-ordinated activities of immune cells, satellite cells, fibroblasts and other resident muscle cells and is controlled by the muscle microenvironment. The sequential steps in the repair pathway are transient and well-orchestrated in normal muscle and tissue homeostasis is generally restored; however, in muscular dystrophy the tissue damage is chronic, inflammatory cells and fibroblasts are continually activated, and satellite cells are less able to proliferate and differentiate to repair the muscle. Adipocytes infiltrate the fibrotic regions compounding the muscle pathology. DMD is associated with chronic inflammation and drugs that reduce inflammation are likely to be beneficial(13-15).

The mouse *mdx* mutation arose spontaneously and is a single base substitution that introduces a premature stop codon in exon 26 of the dystrophin gene (*Dmd*)(16). The *mdx* mouse has no detectable dystrophin protein at the sarcolemma. The phenotype of the *mdx* mouse has been reported in numerous papers and reviews (17-19), so is only briefly reviewed here. Serum creatine kinase increases from around 1 week indicating muscle membrane leakiness, and there is acute onset of myofibre necrosis at 3 weeks identified by the presence of inflammatory cells and degenerating myofibres. Satellite cells (resident muscle progenitor cells) are activated to regenerate the muscle; the newly formed myofibres have centrally located nuclei, rather than peripheral nuclei as in undamaged myofibres. Adult *mdx* mice have a reduced but persistent level of chronic damage and regeneration and the muscle is replaced by ECM (fibrosis) and fatty deposits.

Benfotiamine is a thiamine (vitamin B1) analogue that influences multiple cellular pathways including inhibiting the formation of advanced glycation end products in diabetes (20), reducing inflammation (21), reducing oxidative stress (22) and activating the Akt pathway in heart, endothelial cells and skeletal muscle in diabetic mice (23-26). Benfotiamine is lipid-soluble and can pass through the cell membrane before being converted to biologically active thiamine, increasing levels of thiamine derivatives in the blood and liver, but not in the brain (20, 27-29). Previous research has focused on benfotiamine as a therapeutic for a variety of diabetic related complications including cardiomyopathy (23, 24), retinopathy (30), limb ischaemia (26) and nephropathy (30, 31). It has an excellent safety profile in humans and has been used in many clinical trials without adverse side effects (32-35). Because benfotiamine treatment addresses many of the symptoms associated with muscular dystrophy, we tested benfotiamine treatment in the *mdx* mouse model of Duchenne muscular dystrophy

The current study demonstrates that benfotiamine reduced multiple measures of dystrophic pathology and improved muscle function and performance in the dystrophic *mdx* mouse. Our data suggests these effects could be mediated by reducing inflammation. Due to its excellent safety profile and use in clinical trials for other diseases (31, 32, 36-38), benfotiamine could be transitioned rapidly into a clinical setting, providing benefit to many DMD patients.

## Materials and Methods

All reagents were purchased from Sigma Aldrich (Castle Hill, New South Wales, Australia) unless otherwise specified.

### Animals

All animal experiments were approved by the University of Melbourne Animal Ethics Committee (AEC) and the Murdoch Children’s Research Institute AEC. Mice were purchased from The Animal Resources Centre (Perth, Western Australia) and cared for according to the ‘Australian Code of Practice for the Care of Animals for Scientific Purposes’ published by the National Health and Medical Research Council (NHMRC) Australia (39). They were housed under a 12 hour light/dark cycle with food and water provided *ad libitum*.

### Trial design

Male *mdx* mice were fed a control chow diet, or a diet designed to deliver benfotiamine (Sigma Aldrich) at 10mg/kg bodyweight/day (prepared by Specialty Feeds, Glen Forrest, Western Australia) from 4 weeks of age for 12 weeks. At 10 weeks half the mice, control and benfotiamine trated, were placed into individual cages and allowed access to an exercise wheel; activity was recorded as rotation of the wheel every 1 minute as we have described previously(40). On completion of the trial the mice were anaesthetized with isofluorane, blood was obtained via cardiac puncture and the mice were humanely euthanised via cervical dislocation. Skeletal muscles were harvested and either snap frozen in liquid nitrogen and stored at -80°C for RNA or protein extraction or mounted in 5% tragacanth (w/v) and frozen in liquid nitrogen cooled isopentane and stored at -80°C for histological analyses.

### Immunohistochemistry

Transverse muscle cryosections (10μm) were brought to room temperature and rehydrated in 1xPBS. For immunohistochemical analysis of fibre size, transverse sections were stained for laminin-α2 as follows. Sections were blocked in 10% (v/v) donkey serum (Millipore, Billerica, Massachusetts, USA) in wash buffer (0.1%Tween, 0.5%BSA in 1xPBS) for one hour, and then incubated with the primary antibody, laminin-α2 (Santa Cruz Biotechnology, 1:200), diluted in wash buffer overnight at 4°C. The sections were washed in wash buffer and then incubated with the fluorescent secondary antibody, donkey anti-rat IgG Alexa Fluor 594 (Life Technologies, 1:250) in the dark for 90 minutes. Following a final wash with wash buffer, nuclei were stained with 1μg/μl Hoechst (Life Technologies) in 1xPBS for one minute before mounting with polyvinyl alcohol with glass coverslips. Sections were imaged on a Zeiss Axio Imager M1 upright fluorescent microscope with an AxioCam MRm camera running AxioVision software V4.8.2.0.

For immunohistochemical staining of utrophin associated sarcolemmal proteins the following procedure was used. Avidin/biotin blocking kit (SP-2001; Vector Laboratories) was used according to manufacturer’s instructions. Primary antibodies were prepared using the Mouse on Mouse blocking kit (BMK-2202; Vector Laboratories). Sections were incubated in primary antibody in PBS at 41⍰°C overnight with the following antibodies or lectins: utrophin (MANCHO3; 1:5; Developmental Studies Hybridoma Bank), α-DG (IIH6, sc-53987; 1:500; Santa Cruz Biotechnology), β-DG (VP-B205; 1:50; Vector Laboratories), α-SG (VP-A105; 1:30; Vector Laboratories), β-SG (VP-B206; 1:30; Vector Laboratories), γ-SG (VP-G803, 1:100; Vector Labs), SSPN (E-2; 1:100; Santa Cruz Biotechnology), and WFA (B-1355; 1:500; Vector Laboratories). Primary antibodies were detected with a biotinylated anti–mouse IgG antibody (BA-9200; 1:500; Vector Laboratories). Fluorescein–conjugated avidin D (A-2001; 1:500; Vector Laboratories) was used to detect secondary antibodies and biotinylated WFA. Sections were mounted in Vectashield (Vector Laboratories) and visualized using an Axioplan 2 fluorescence microscope with Axiovision 3.0 software (Zeiss). Images were captured under identical conditions.

### Histology

Hematoxylin and eosin (H&E) staining was used to visualise the structure of the muscle myofibres, nuclei and connective tissue. Entire H&E stained sections were imaged on a Mirax Scan bright field automated digital slide Scanner (Carl Zeiss, Oberkochen, Germany) at the University of Melbourne Department of Anatomy and Neuroscience (http://www.apn-histopathology.unimelb.edu.au/).

### Measurements of pathology

Images were analysed using Image J version 1.48G (U. S. National Institutes of Health, Bethesda, Maryland, USA). Myofibre diameter was measured as Minimum Feret’s diameter from laminin-α2 stained transverse sections as per Treat NMD guidelines (http://www.treat-nmd.eu/downloads/file/sops/dmd/MDX/DMD_M.1.2.001.pdf). Damaged myofibres were detected using IgG staining, and the percentage of IgG positive myofibres from the entire quadriceps cross section was calculated. Areas of damaged tissue were defined based on the presence of infiltrating inflammatory cells and areas of degenerating myofibres with fragmented sarcoplasm by hematoxylin and eosin staining as per the Treat-NMD standard operating procedure (http://www.treat-nmd.eu/downloads/file/sops/dmd/MDX/DMD_M.1.2.007.pdf). The extent of tissue damage was expressed as a percentage of the total quadriceps area. Central nucleation was expressed as the percentage of centrally nucleated myofibres over the total muscle cross-section.

### Creatine kinase assay

Blood obtained from cardiac puncture was centrifuged at 12,000g for 15 minutes to separate the serum from the other components. Serum was aliquoted into sterile tubes and stored at -80°C until required. The cell lysates were thawed and 5μl of each lysate was mixed in triplicate with 100μl of CK-NAC (Thermo Scientific, Waltham, Massachusetts, USA). The change in absorbance was recorded at 340nm over three minutes (measured in 20 second intervals) at 37°C using a Paradigm Detection Platform (Beckman Coulter, Brea, California, USA).

### RNA extraction, cDNA synthesis and qPCR

RNA was extracted with TriReagent (Sigma Aldrich) followed by purification and DNAse treatment using the SV Total RNA Isolation System (Promega). cDNA was synthesised from 1μg total cellular RNA with MML-V Reverse Transcriptase (Promega). Gene expression was quantitated using qPCR as previously described (41).

Oligonucleotide sequences are presented in Table 1. Primers were designed using ‘Primer-BLAST’ available on the website http://www.ncbi.nlm.nih.gov, Primers were tested for efficiency by serial dilution of cDNAs. All primers had an efficiency between 1.8 and 2.2 per cycle. Data are expressed as the mean of normalised expression to the housekeeper hypoxanthine-guanine phosphoribosyltransferase (*Hprt*) (41).

**Table 1.**
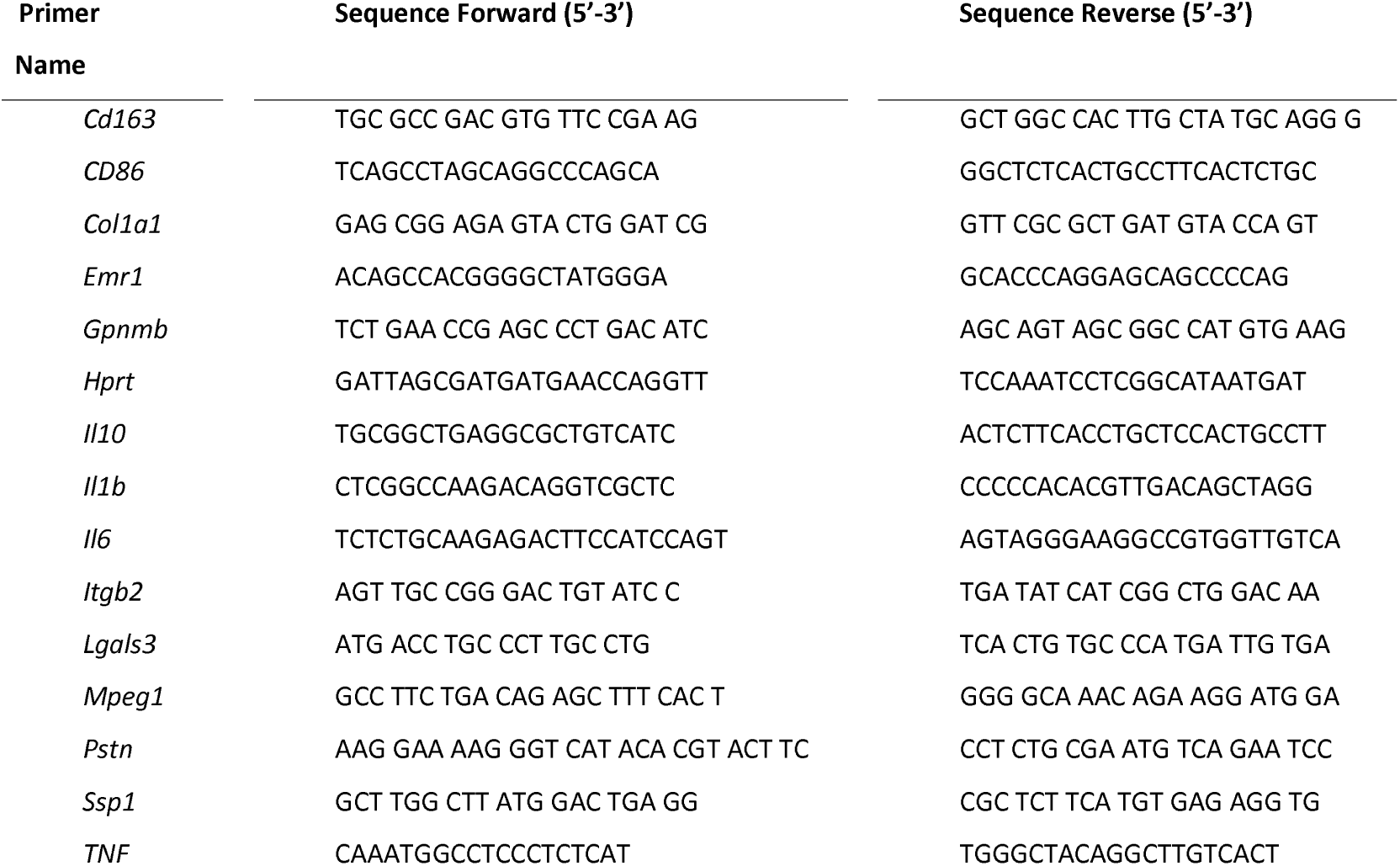
Primer sequences used for qPCR.

### Immunoblots

Protein was extracted from frozen *quadriceps* muscles by homogenization in ice cold extraction protein extraction buffer (1% NP40 alternative, 1mM EDTA, Complete EDTA-free protease inhibitor tablets and PhosSTOP Phosphatase Inhibitor Cocktail Tablets (Roche) in 1xPBS). Protein concentrations were determined using a 2D quant assay (GE Healthcare BioSciences). Protein samples (100μg) were resolved through 4-12% gradient SDS-PAGE (Life Technologies) and transferred to nitrocellulose membranes. The membranes were probed with antibodies to the following proteins,: Akt (Cell Signalling Technologies), phosphorylated Akt Ser 473 (Cell Signalling Technologies), α-tubulin (Sigma Aldrich), followed by HRP-conjugated anti-mouse IgG (Cell Signalling Technologies). Detected was performed using Amersham ECL Western Blotting reagents and relative protein levels were quantitated using an ImageQuant LAS400 and Image Quant TL 7.0 software (GE Healthcare and BioSciences).

### Forelimb grip strength measurements

Forelimb grip strength was measured weekly from 4 weeks of age using a BIO-GS3 grip strength meter (Bioseb In Vivo Research Instruments). The procedure was performed as per the Treat-NMD standard operating procedure (http://www.treat-nmd.eu/downloads/file/sops/sma/SMA_M.2.1.002.pdf). The force of the pull (N) prior to release was recorded and the mouse was placed back in its cage and allowed to recover for 5-9 minutes before repeating the test another 4 times.

### Statistical analyses

Where there was a direct comparison between two groups or two data points, a Student’s t-test was used to compare the mean and standard error of the data. The statistical package Genstat (14th Edition) was used to compare multiple groups. A one-way analysis of variance (ANOVA), with Dunnet’s or Bonferroni post-hoc test was used when all treatments were compared to a single control or all treatments compared to each other respectively. Analysis that included time as a variable, grip strength, was conducted using a two-way ANOVA .qPCR data was assessed using a non-parametric Mann-Whitney U test in GraphPad as described in Pfaffl (42).

## Results

### Benfotiamine increases growth and promotes myofibre hypertrophy in mdx mice

Body weights of the three experimental groups, benfotiamine treated *mdx*, control *mdx* and wild type were compared over the treatment period (Figure 1A). Benfotiamine treated *mdx* mice were heavier than control *mdx* mice in the initial three weeks of treatment (ages 4 to 7 weeks) corresponding to the period of rapid growth. Body weights did not differ between groups during the middle phases of the treatment period. In the last 2 weeks bodyweights were similar in control *mdx* and wild type but benfotiamine treated mice were heavier. The last two weeks were the only time when the body weight of benfotiamine treated *mdx* mice was significantly different from wild type mice. We also weighed quadriceps, tibialis anterior, gastrocnemius, extensor digitorum longus, soleus and heart muscles; when corrected for body weight, there were no significant differences between groups (data not shown).

**Figure 1.**
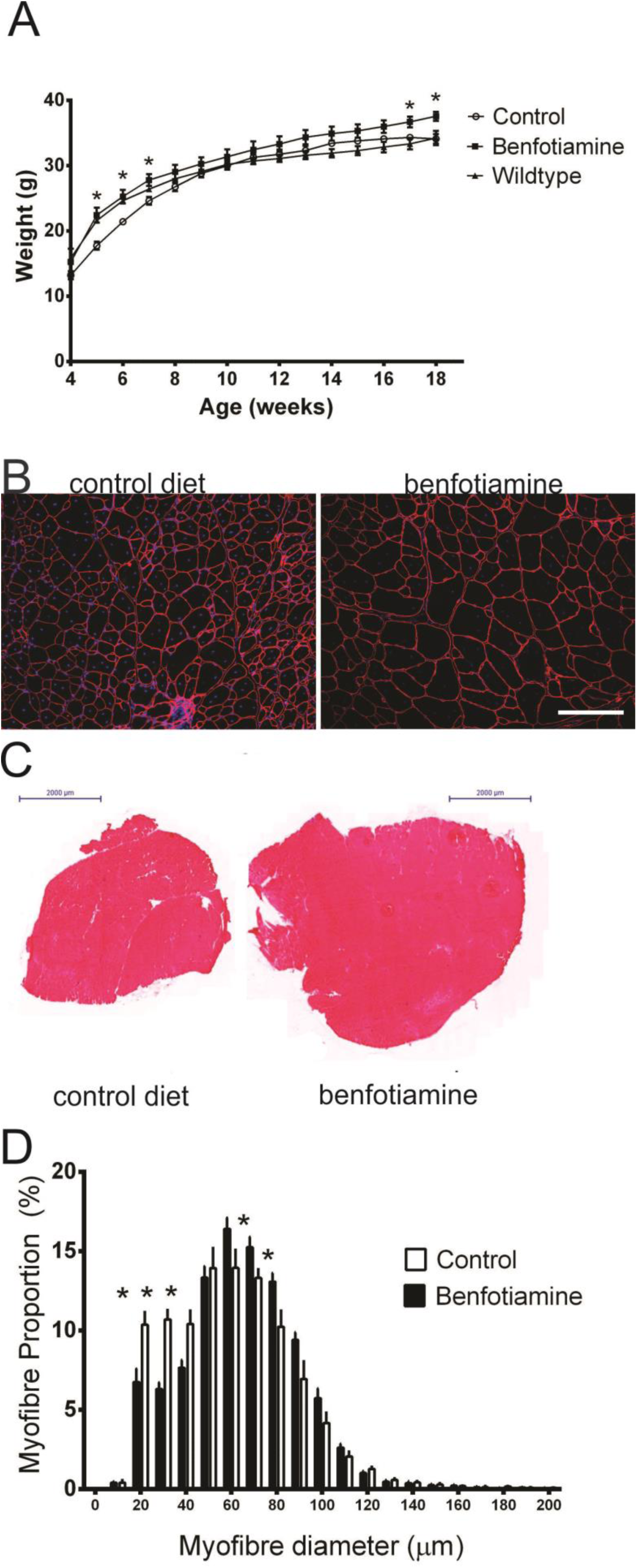
Benfotiamine increases *mdx* body weight and quadriceps myofibre diameter. (A) Mice were weighed weekly for the 12 weeks of the trial. Benfotiamine treated *mdx* mice weighed more than the *mdx* control cohort from 5-7 weeks of age, then from 17-19 weeks of age. Benfotiamine treated *mdx* mice were similar in weight to wild type mice across the entire treatment period. The quadriceps muscle from control (B) and benfotiamine treated (C) *mdx* mice were stained with a laminin α2 antibody (red) and DAPI (blue). Benfotiamine increased the quadriceps cross sectional area by increasing the number of larger (>70μm diameter) myofibres and decreasing the number of smaller (<40μm diameter) myofibres (D). The graphs show mean ± s.e.m. * indicates p<0.05 (benfotiamine compared to *mdx* control), n=6. Scale bar, 200 μM.

To determine the effect of benfotiamine on muscle fibre diameter, we examined the Feret’s minimal diameter of muscle fibres in the quadriceps from control and benfotiamine treated *mdx* mice Figure 1B,C. There was a significant decrease in the proportion of smaller myofibres (10-30µm) in the quadriceps from the benfotiamine treated *mdx* mice (Figure 1D) (p<0.05). Conversely, there were more larger myofibres (70-80µm) in the *mdx* benfotiamine treated quadriceps (Figure 1D)(p<0.05).

### Benfotiamine reduces markers of dystrophic pathology

Dystrophic pathology in patients and *mdx* mice is characterised by elevated serum creatine kinase, myofibre degeneration, immune cell infiltration and replacement of muscle with connective tissue. Transverse *quadriceps* sections were stained with IgG to mark myofibres with compromised sarcolemmal integrity (Figure 2A and B). The percentage of IgG positive myofibres was reduced in the *quadriceps* of benfotiamine treated *mdx* mice when compared to the *mdx* mice on the control diet ((p<0.05) Figure 2C). Skeletal muscle damage – areas with infiltrating inflammatory cells and degenerating myofibres - was reduced by approximately 48% in benfotiamine treated *mdx* mice, when compared to the *mdx* mice on the control diet ((p<0.05) Figure 2D). Serum creatine kinase (CK), indicative of “leaky”/damaged muscle fibres, was reduced by around 34% inbenfotiamine treated *mdx* compared to control *mdx* mice (p=0.073) (Figure 2E).

**Figure 2.**
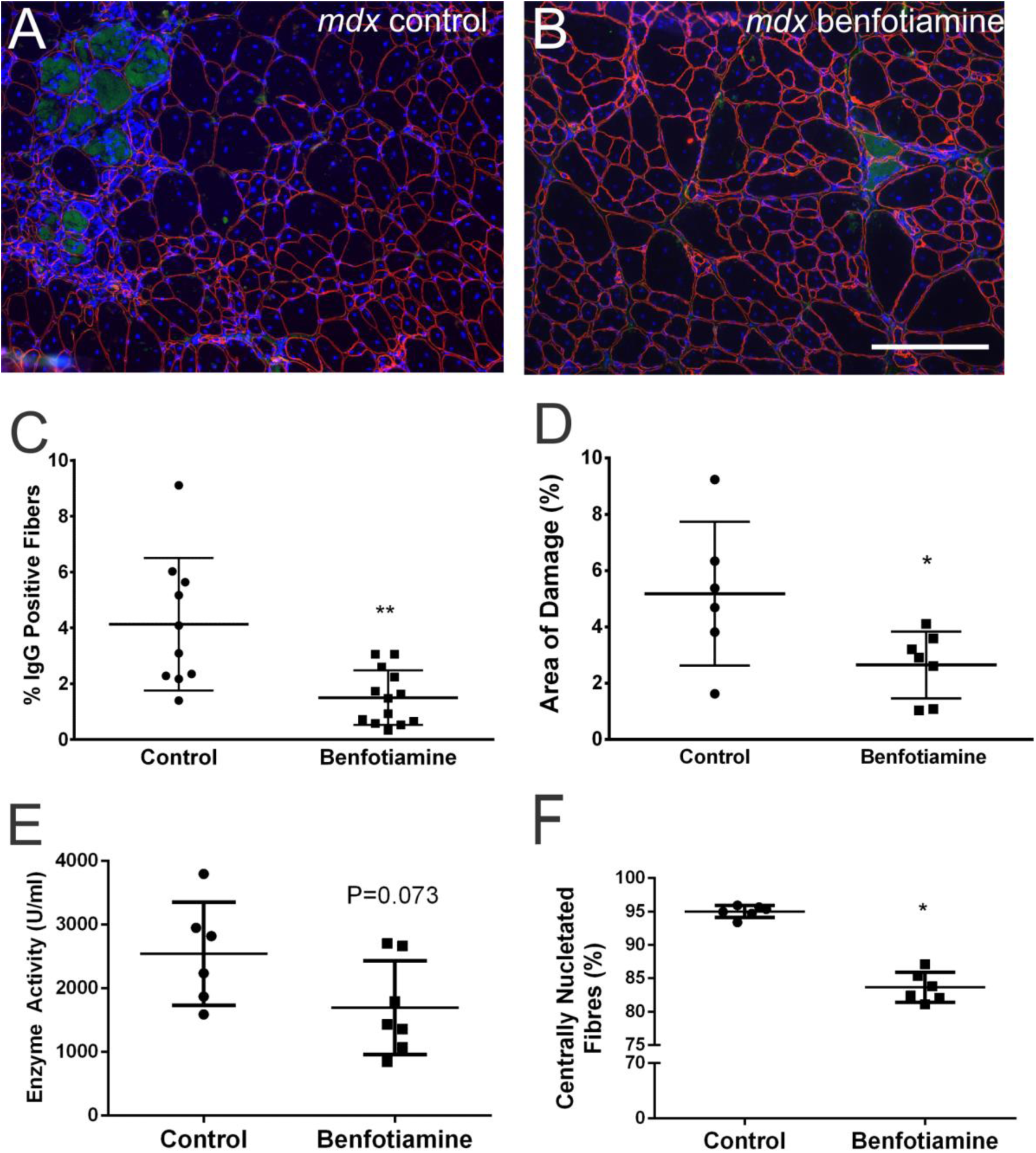
Benfotiamine reduces muscle damage and improves sarcolemma stability in *mdx* mice. Transverse quadriceps cryosections from *mdx* mice on control diet (A) or benfotiamine (B) co-stained with antibodies to laminin α2 antibody (red) and IgG (green). Nuclei are stained with DAPI (blue). Benfotiamine significantly reduced the percentage of damaged myofibres permeable to IgG (C). Benfotiamine reduces the area of damage (including areas of necrosis and inflammatory cell infiltration) observed in H&E stained transverse sections compared to *mdx* mice on a control diet (D). Muscle-specific creatine kinase in the serum, a marker of sarcolemma damage, is reduced with benfotiamine treatment (E). Benfotiamine also reduces the proportion of fibres in mdx skeletal muscle with central nucleation (F). *p<0.05, **p<0.001, n=6. Scale bar, 200 μM.

The reduction in dystrophic pathology seen with benfotiamine administration could occur by promoting growth of new myofibres, or by preventing skeletal muscle damage. To investigate these mechanisms the percentage of fibres with central nucleation was measured over the quadriceps cross section. In healthy muscle, the nuclei are located peripherally, beneath the basal lamina, but the nuclei of regenerated muscle fibres are centrally located. Benfotiamine administration significantly reduced the percentage of myofibres with centrally located nuclei in the *mdx* quadriceps ((p<0.05), Figure 2F). This finding suggests that the primary effect of benfotiamine is to protect the muscle from damage rather than to increase regeneration.

### Forelimb grip strength is increased in benfotiamine treated mdx mice

To determine if the reduced dystrophic pathology observed with benfotiamine treatment translates to functional improvements in strength and performance, forelimb grip strength was measured. Wildtype mice were consistently stronger (61-107%) than *mdx* mice on the control and benfotiamine diets across the entire treatment period (Figure 3A). Benfotiamine treated *mdx* mice were stronger than the *mdx* control cohort from 4 weeks of treatment (p<0.01) until trial completion (p<0.0001) (Figure 3A)..

**Figure 3.**
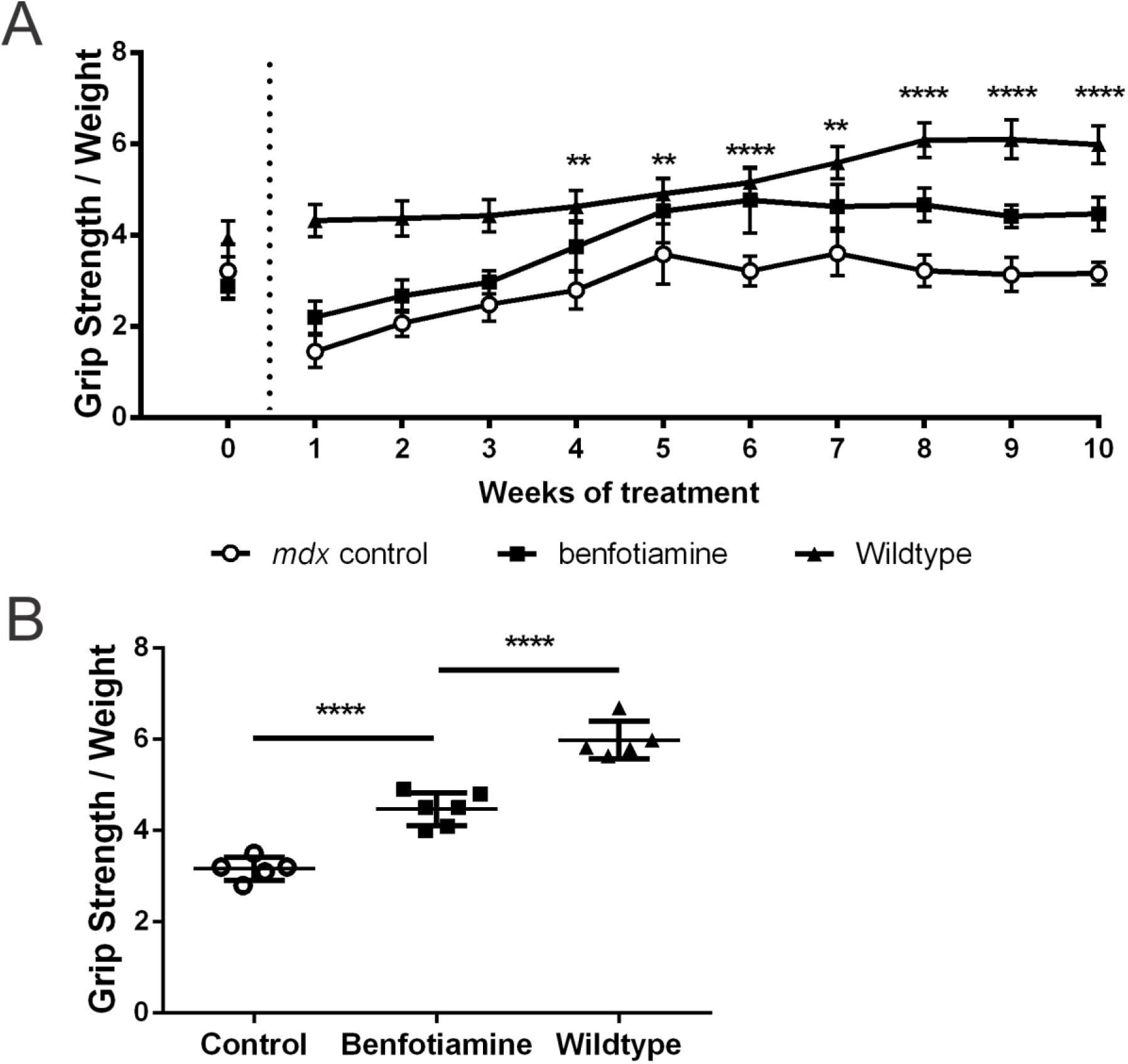
Benfotiamine increases grip strength in *mdx* mice. (A) Forelimb grip strength was measured in benfotiamine treated, control *mdx* and wild type mice on a weekly basis. Wild type mice were stronger than both *mdx* groups throughout the trial (p<0.0001 with two-way ANOVA). After 4 weeks of treatment grip strength was significantly increased in benfotiamine treated *mdx* mice compared to *mdx* controls; this difference was maintained till the end of the trial. (B) At the completion of the treatment grip strength in benfotiamine treated *mdx* mice was significantly improved but did not reach wild type levels. Grip strength was normalised to bodyweight. **p<0.01, ***p<0.001, ****p<0.0001 indicates differences between mdx mice on control and benfotiamine diets, n=6. Graphs show mean ± SEM. (n=6).

At the end of the trial, grip strength was increased in the benfotiamine treated *mdx* group compared to control *mdx* mice ((p<0.001), Figure 3B), but had not reached wild type levels. These data reveal that benfotiamine is an effective treatment to reduce pathology and improve muscle function in *mdx* mice.

### Benfotiamine treatment improves voluntary exercise performance

Voluntary exercise was recorded during the last two weeks of the trial and exercise parameters including the running time and distance and speed, rest time, and the number of run bouts were calculated (Figure 4). Overall benfotiamine treated *mdx* mice ran further and faster than their control counterparts. The daily mean distance was increased compared to controls (p<0.05) as was the total distance that treated mice ran over the trial period (Figure 4A). There was no difference in the mean rest time between groups (Figure 4B); benfotiamine treated *mdx* rested for a shorter time in total indicating that they rested less often. There was no difference between the treatment groups in the number of run bouts (Figure 4C), but benfotiamine treated mice ran for longer (Figure 4D) and covered more distance per exercise bout (p<0.05) (Figure 4E). The rate at which benfotiamine treated mice ran was higher than controls (p<0.05) (Figure 4F). There was no difference in the maximum run rate (maximum speed) with benfotiamine treatment (not shown).

**Figure 4.**
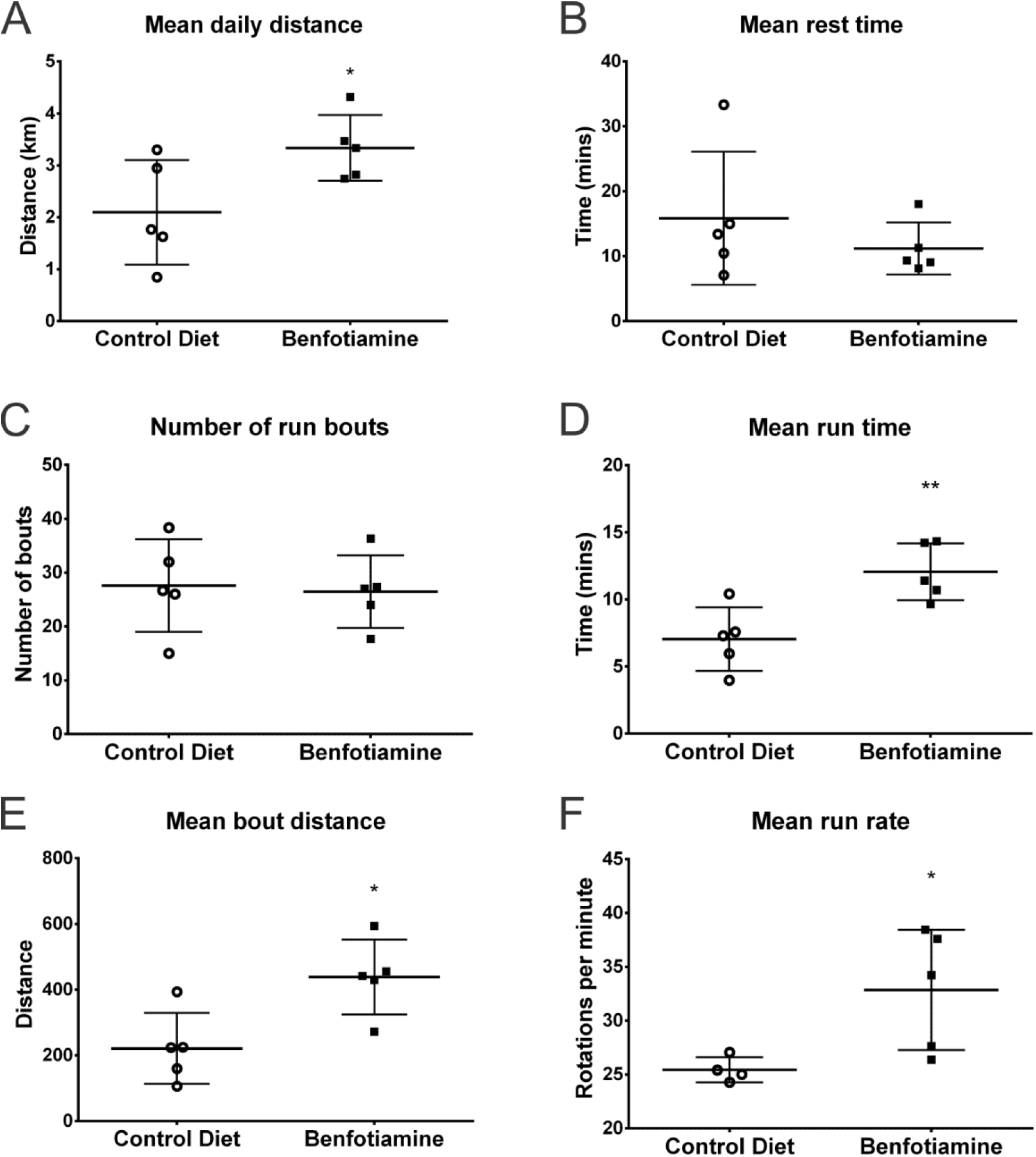
Benfotiamine improves voluntary exercise performance. For the final 2 weeks of the trial mice were allowed access to an exercise wheel. A number of parameters were recorded or calculated to examine the effect of benfotiamine on exercise performance. Benfotiamine treated mice ran further each day (A) but rested for similar periods of time (B) and ran on the exercise wheel a similar number of times each day (C). Benfotiamine treated mice ran for a longer period of time (D), covered a greater distance each time they ran (E) and ran at a faster rate (F) than did control mice. The graphs show mean ± s.e.m. * indicates p<0.05, n=6. Scale bar.

### Benfotiamine treatment increases compensatory cell-matrix adhesion complexes

Protection from myofibre damage in dystrophic skeletal muscle has been associated with increased expression of adhesion complexes, including components of the utrophin glycoprotein complex (UGC) and α7β1 integrin, which is thought to compensate for the lack of dystrophin (43, 44). During fetal development, utrophin is found around the entire sarcolemma of developing myofibres; however, in adult skeletal muscle utrophin is restricted to the post-synaptic region of the neuromuscular junction (45, 46). Increased utrophin compensates for dystrophin loss, and associates with β-dystroglycan. In turn, this is further stabilized by the sarcoglycan-sarcospan sub-complex (consisting of α-, β-γ- and δ-sarcoglycan)(44). Given the histopathological improvements in benfotiamine-treated *mdx* tissue, we examined the abundance and distribution of utrophin and associated UGC proteins. Expression of utrophin (*Utr*) was not increased with benfotiamine (not shown); however, immunohistochemistry demonstrated that benfotiamine treatment increased UGC protein staining at the sarcolemma in *mdx* muscle (Figure 5).

**Figure 5.**
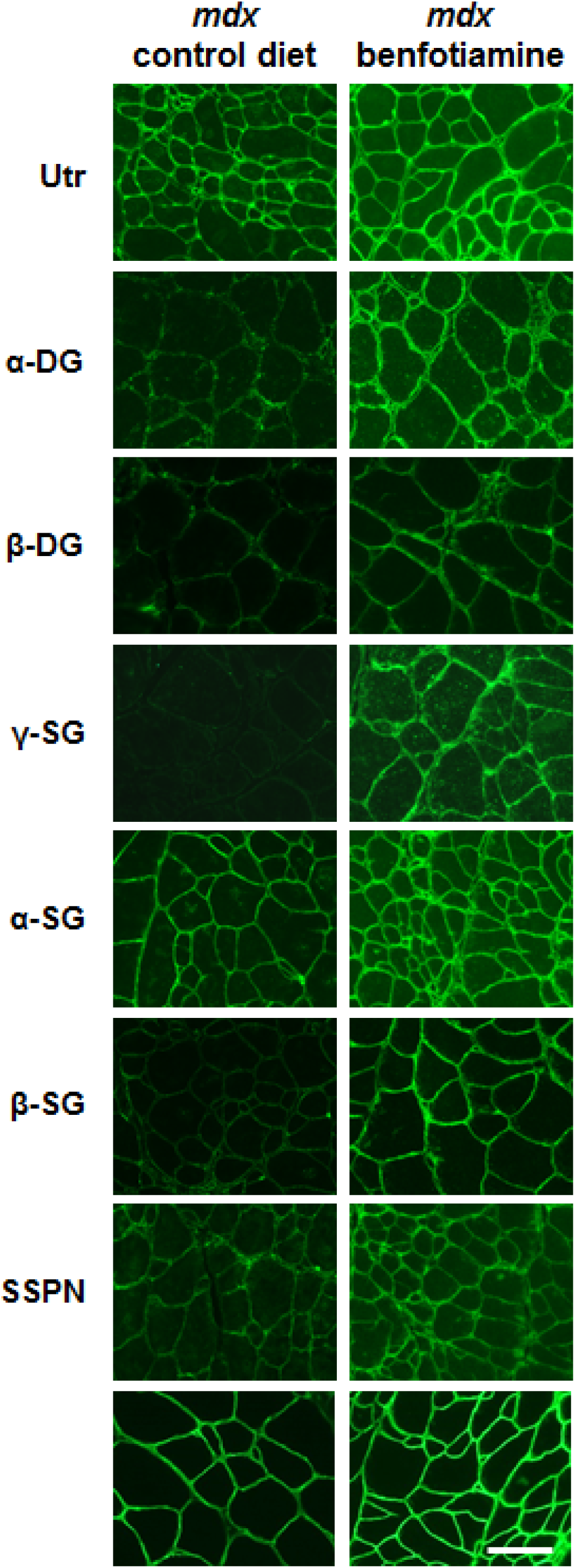
Benfotiamine treatment increases localisation of utrophin-glycoprotein complex to the sarcolemma. Immunofluorescence analysis of benfotiamine treated and control *mdx* quadriceps reveals an increased abundance of utrophin-glycoprotein complex at the sarcolemma. Utrophin (utrn), α-/β-dystroglycan (α-/β-DG), α-/γ-/β-sarcoglycan (α-/-γ/β-SG), sarcospan (SSPN) are shown. Bottom panels are stained with wheat germ agglutinin (WFA) to define the sarcolemma. Scale bar, 50 μm.

### Benfotiamine does not activate Akt signaling in *mdx* skeletal muscle

*Akt* and *mTOR* mRNAs were significantly up regulated in muscle from *mdx* mice fed the benfotiamine diet (data not shown); however, total Akt protein expression was not changed by benfotiamine, and the proportion of phosphorylated Akt (the active form; p-Akt) following benfotiamine treatment was not increased (data not shown). Levels of mTOR and phosphorylated mTOR were also not changed in muscle from treated mice (not shown).

### Benfotiamine reduces gene expression of inflammatory markers

Dystrophic pathology progression is associated with chronic inflammation that disrupts normal homeostasis and exacerbates pathology by up-regulating inflammatory cytokines, chemokines and immune cells (47-51). To determine if benfotiamine has anti-inflammatory effects in the context of *mdx* skeletal muscle, we analysed the relative gene expression of macrophage markers and pro-inflammatory cytokines. Expression of the pan-macrophage marker *Emr1* was downregulated with benfotiamine treatment in the *mdx* mice (p<0.05) (Figure 6A). *Cd86* (a marker of cytotoxic M1 macrophages) was also down-regulated (p<0.05) (Figure 6B) but *Cd163* mRNA (M2 macrophages) was unchanged with benfotiamine treatment (not shown). The pro-inflammatory cytokines, *Tnf* and *Il1b*, were reduced in the *mdx* mice with benfotiamine treatment when compared to the control *mdx* mice (p<0.05) (Figure 6C and D); however, *Il6* and *Il10* mRNAs were unchanged (Figure 6E and F).

**Figure 6.**
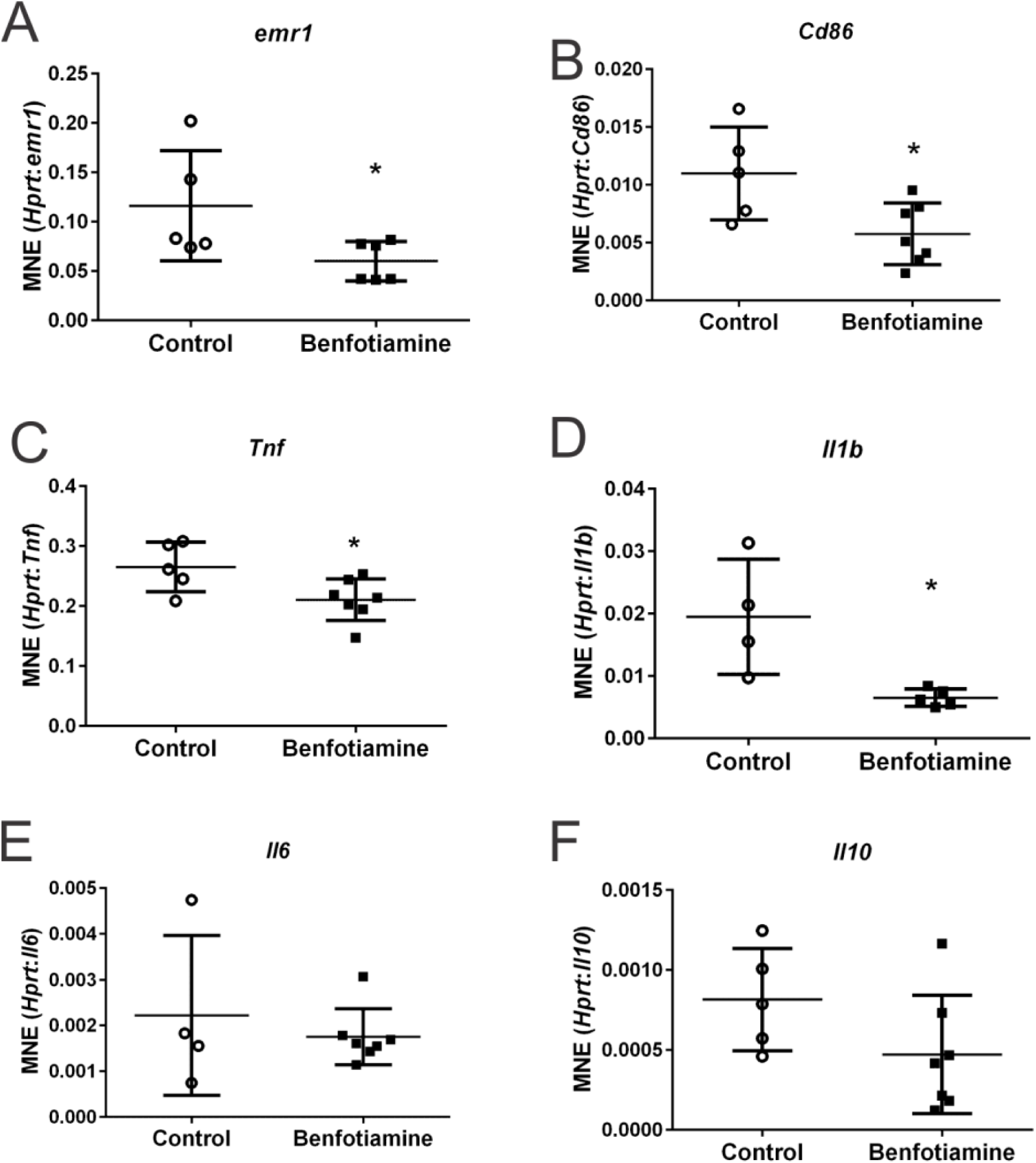
Benfotiamine administration down-regulates expression of pro-inflammatory cytokines and markers of macrophages in treated *mdx* mice. The pan macrophage marker Emr1 (A) and M1 macrophage marker *Cd86* (B) mRNAs were down-regulated with benfotiamine administration compared to *mdx* controls. *Cd163* (a marker for M2 macrophages) expression was unchanged (not shown). *Tnf* (C) and *IL1b* (D) mRNAs are down-regulated, and both *Il6* (E) and *Il10* (F) are unchanged with benfotiamine treatment. Graphs show mean ±SEM. * indicates p<0.05 (n=5-7).

Previous transcript profiling studies and our own unpublished data showed altered expression of pathways involved in the inflammatory/immune response in *mdx* mice(52-54). Based on these studies, we examined expression of a number of inflammatory genes that have previously been reported as the most relevant differentially expressed genes; *Mmp12, Gpnmb, Postn, Mpeg1, Lgals3, Itgb2* and *Spp1* (Figure 7A-G). Of these genes, only *Itgpb2* expression was not affected by benfotiamine treatment. *Spp1, Postn*, and *Col1a1* are all strongly up-regulated in dystrophic skeletal muscle and are down-regulated with benfotiamine treatment (Figure 7C, G and H).

**Figure 7.**
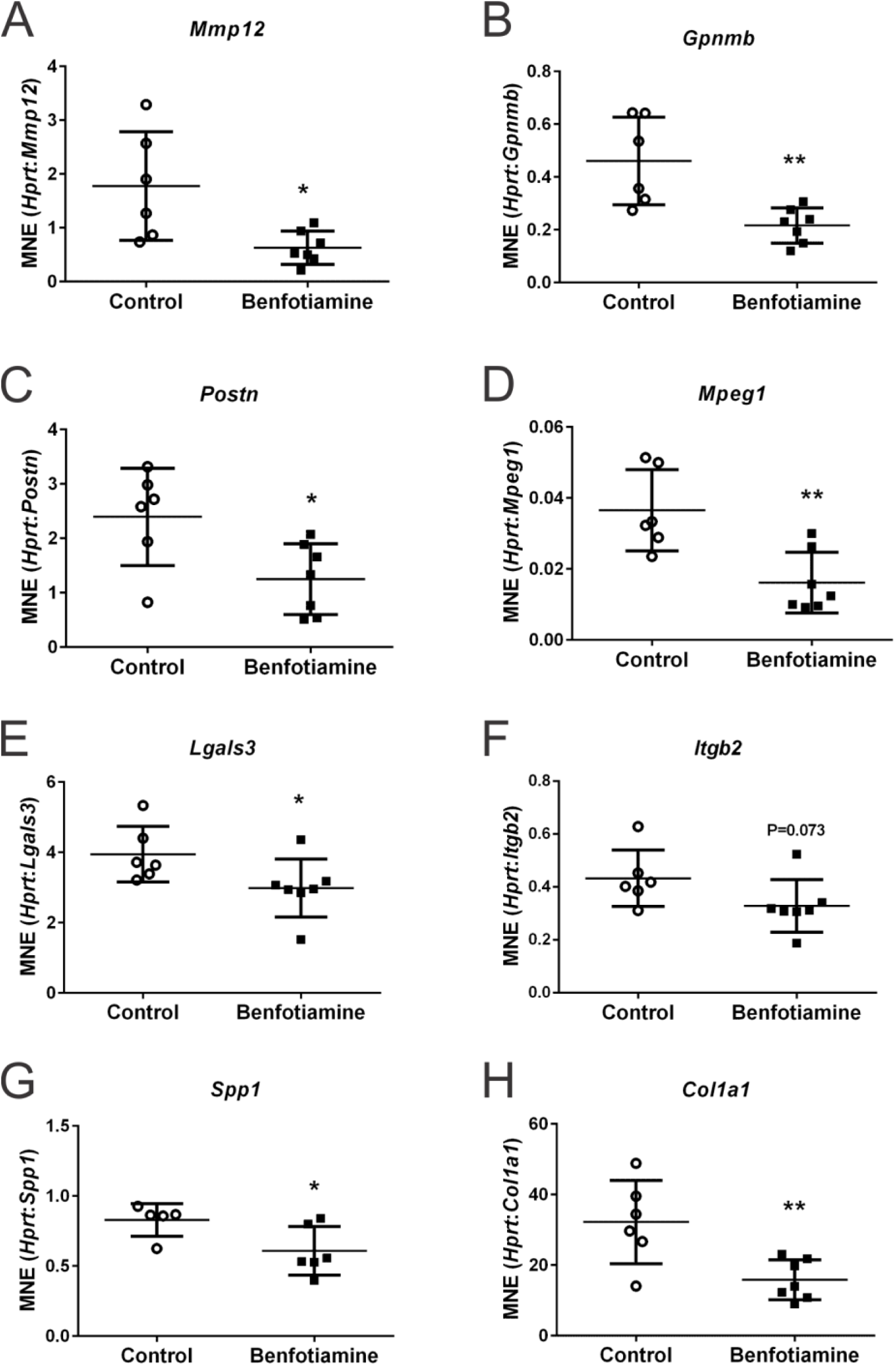
Expression of genes associated with inflammatory responses and fibrosis are reduced by benfotiamine treated in *mdx* mice. *Mmp12* (A), *Gpnmb* (B), *Postn* (C), *Mpeg1* and *Lgals3* (E) mRNAs were all down regulated by benfotiamine treatment. *Itgb2* (F) expression was decreased following treatment but this was not significant (P=0.073). *Spp1* (G) and *Col1a1* (H) mRNAs were also decreased compared to controls in *mdx* muscle following treatment. Graphs show mean ±SEM. * indicates p<0.05 ** indicates P<0.01 (n=5-7).

## Discussion

In this study, we have shown that benfotiamine, a lipid soluble analogue of vitamin B1, reduces multiple measures of dystrophic pathology and improves muscle strength and performance in *mdx* mice. The histopathological changes associated with disease progression in DMD include many fibres with central nucleation, wider variation in fibre diameter, extensive necrosis, chronic inflammation, fat deposition and tissue fibrosis (13, 55, 56). The progressive accumulation of collagen and related ECM proteins and the apparent dysregulation of matricellular proteins are believed to play an important role in DMD, with the progressive loss of muscle fibres and their replacement with non-contractile fibrotic tissue being a major histopathological hallmark that correlates with reduced motor function (57). In fact, the excessive accumulation of ECM components is an indicator of the decline in muscle strength (58).

We originally hypothesised that benfotiamine treatment would increase phosphorylation of the components of the AKT signalling pathway and this in turn would increase utrophin expression resulting in protection from damage(43). Benfotiamine clearly protects *mdx* skeletal muscle from damage, but we did not see any significant effect on the AKT signalling axis. Utrophin expression is increased in *mdx* muscles (45, 59) and is more abundant around the sarcolemma of regenerating fibres than in mature fibres (60). In situations where utrophin is upregulated through transgenic or pharmacological means, dystrophic muscle fibres are protected from damage (61, 62). While we do not observe utrophin upregulation at the transcript level in response to benfotiamine, we do find evidence of increased localisation of utrophin around muscle fibres.

Inflammation is a major contributor to disease progression in DMD and one of the major components of this response is the infiltration and activity of macrophage populations. We show a decrease in both the pan macrophage marker *Emr1* and the M1 macrophage marker *Cd86* in benfotiamine treated muscle and a decrease in pro-inflammatory *Tnf* and *Il1b* gene expression. We also provide evidence for the down regulation of other genes associated with macrophage activity and function.

MMP12, often referred to as macrophage elastase, is primarily produced by macrophages and is associated with a number of pathological conditions including aortic aneurism, atherosclerosis, emphysema and rheumatoid arthritis (63). MMP12 activity is elevated during ECM remodelling but it also cleaves non-ECM targets such as latent TNF (64). As well as activating TNF, it has pro-inflammatory activity that can recruit neutrophils and increase cytokine and chemokine production. MMP12 can cleave and activate chemokines such as mCXCL5, hCXCL5, and hCXCL8 which are involved in recruiting neutrophils. Macrophages accumulate at injury sites after 24-48 hours and secrete MMP12 to inactivate these same chemokines and contribute to the reduction in neutrophils at the site of damage. MMP12 can also inactivate CCL2, -7, -8, and -13 further assisting to resolve the inflammation (63).

A number of transcriptomic studies have examined the gene expression patterns that underlie DMD progression. The method of analysis used to identify dysregulated pathways is not consistent between these studies (65) but a common theme is a chronic inflammatory response(54). Clusters of genes which contribute to progression including *Spp1, Itgbp2, Mpeg1, Postn, lgals3, Gpnmb* and *Mmp12*(*53*) are also often upregulated. Interestingly studies that show protection from damage in *mdx* muscles mediated by increased utrophin staining at the sarcolemma also report an effect on this gene network (52). Benfotiamine treatment of *mdx* mice resulted in down regulation of all these genes, except *Itgb2*, that have previously been associated with disease progression. The changes we report here in gene expression after benfotiamine treatment are consistent with previous studies, some in other tissue systems, which indicate a dampening of the pro-inflammatory response. Glycoprotein nonmetastatic melanoma protein B (GPNMB) is involved in inflammation and fibrosis after tissue injury. The expression of *Gpnmb* has been associated with increased damage in *mdx* muscle (66) and there is significant evidence that GPNMB is associated with inflammatory disease of cardiac muscle. GPNMB adversely influences myocardial remodelling(67) and *Mmp12* and Gpnmb expression have been associated with inflammatory processes associated with myocarditis(68).

Osteopontin (*Spp1*) is a matricellular protein that is increased along with periostin (*Pstn*) in muscular dystrophy (53, 54, 69, 70). Osteopontin is highly expressed in dystrophic muscle, is associated with the inflammatory infiltrate during regeneration and is closely linked to fibrosis in skeletal muscle (72). In osteopontin and dystrophin double knockout mice, this matricellular protein acts as an immune-modulator in skeletal muscle and a pro-fibrotic cytokine in muscular dystrophy (73). Dysregulation of both osteopontin and periostin is an early feature in laminin-deficient muscular dystrophy (74, 75); indicating that changed levels of both these matricellular proteins are key factors involved in development of muscular dystrophy related fibrosis(76). These studies clearly establish matricellular proteins as promising therapeutic targets for reducing fibrosis in muscular dystrophy. Agents such as benfotiamine which correct secondary abnormalities in ECM protein expression, including matricellular proteins, could reduce scar tissue accumulation, and maintain skeletal muscle elasticity and function.

Fibrosis is pronounced in DMD and the *mdx* mouse model, and is a major contributor to muscle dysfunction and disability. The *mdx* mouse shows varying degrees of fibrosis (77, 78) with increased collagen and proteoglycan expression seen in dystrophin deficient muscle including the limb muscles and the diaphragm (79-81). A recent proteomic characterisation of the diaphragm in *mdx*-4cv, an *mdx* genetic variant which shows decreased revertant fibres, showed that the most significantly increased protein was the matricellular protein periostin (82); this confirms a previous study which showed increased periostin transcript in limb muscles of the same mouse model (83). These studies are in keeping with observations of increased periostin in DMD biopsy material (84). There is a close link between inflammatory signalling, through IL-17, periostin expression and fibrotic effects as exemplified by observations that IL-17 and TNF exert a synergistic effect on the increased expression of type 1 collagen in liver fibrosis (85). Periostin ablation in mice reduces the dystrophic symptoms(84). While we did not directly examine fibrosis in this study there are a number of robust changes in gene expression which suggest that benfotiamine could be reducing fibrosis; *Spp1, Pstn*, and *Col1a1* are all downregulated following benfotiamine treatment.

This is the first study to assess the potential of benfotiamine as a treatment for neuromuscular conditions such as DMD. Further studies can assess whether benfotiamine would provide additive benefits when used with corticosteroids. With many promising treatments for DMD still in development, future studies could allow benfotiamine, with its excellent safety profile and encouraging results in the *mdx* mouse model, to rapidly transition into the clinic, providing benefit to many DMD patients in the interim.

## Acknowledgements

This work was supported by Muscular Dystrophy Australia, Murdoch Children’s Research Institute and the Victorian Government’s Operational Infrastructure Support Program. This work was supported by grants from the National Institutes of Health [R01 AR048179 to R.C.W, T32 AR059033 and F32 AR069469 to E.M.G] and the Muscular Dystrophy Association USA [274143 and 416364 to R.C.W.]. SRL was supported by a National Health and Medical Research Council of Australia research fellowship [GNT1043837].

